# Spatial Heterogeneity of Delta-like 4 Within a Multinucleated Niche Cell Maintains Muscle Stem Cell Diversity

**DOI:** 10.1101/2020.10.20.347484

**Authors:** Susan Eliazer, Xuefeng Sun, Andrew S. Brack

## Abstract

The quiescent muscle stem cell (QSC) pool is heterogeneous and generally characterized by the presence and levels of intrinsic myogenic transcription factors. Whether extrinsic factors maintain the diversity of states across the QSC pool remains unknown. The muscle fiber is a multinucleated syncytium that serves as a niche to QSCs, raising the possibility that the muscle fiber regulates the diversity of states across the QSC pool. Here we show that the muscle fiber maintains a continuum of quiescent states, through a gradient of Notch ligand, Dll4, produced by the fiber and captured by QSCs. The abundance of Dll4 captured by the QSC correlates with levels of the SC identity gene, Pax7. Niche-specific loss of Dll4 decreases QSC diversity and shifts the continuum, towards more proliferative and committed states. We reveal that fiber-derived Mindbomb1 (Mib1), an E3 ubiquitin ligase activates Dll4 and controls the spatial localization of Dll4. In response to injury, with a Dll4-replenished niche, the normal continuum and diversity of SC pool is restored, demonstrating bi-directionality within the SC continuum. Our data shows that a post-translational mechanism controls spatial heterogeneity of Notch ligands in a multinucleated niche cell to maintain a continuum of diverse states within the SC pool during tissue homeostasis.

## Introduction

Adult muscle stem cells (SCs) are essential for muscle tissue repair. Subsets of the SC pool are endowed with self-renewal potential and others are restricted to differentiation (Chakkalakal et al., 2014; Kuang et al., 2007; Rocheteau et al., 2012; Scaramozza et al., 2019). This is consistent with a unidirectional and hierarchical relationship between stem cells. Intrinsic and extrinsic cues regulate cell fate decisions of activated SCs to self-renew or differentiate (Dumont et al., 2015; Kuang et al., 2008). Due to low muscle tissue turnover, SCs exist predominantly in a quiescent state for the majority of adult life. Through single cell RNA-sequencing (scRNA-seq) and transgenic reporter mice it is appreciated that the QSC pool is molecularly and phenotypically heterogeneous (Chakkalakal et al., 2014; De Micheli et al., 2020; Dell’Orso et al., 2019; Kimmel et al., 2020; Kuang et al., 2007; Rocheteau et al., 2012). These different cell states enable SCs to exhibit functional heterogeneity in response to activation and injury cues. It is not known how the diversity of states are maintained across the QSC pool. SC quiescence is actively maintained by paracrine-acting cues from the muscle fiber, serving as a niche cell (Bischoff, 1990; Eliazer et al., 2019; Goel et al., 2017). In contrast to niche cells across many stem cell compartments, each muscle fiber is a multi-nucleated syncytium, that exhibits transcriptional diversity across the myonuclei, to provide spatial control for specialized functions (Kim et al., 2020; Petrany et al., 2020). Does the multinucleated niche cell regulate the diversity of states across the QSC pool?

## Results

### Adult QSCs exist in a continuum of molecular cell states

To answer this question, we first asked whether the QSC pool is composed of a continuum of cell states during tissue homeostasis. We stained freshly isolated single muscle fibers from adult mice, with the SC identify marker, Pax7 (Lepper et al., 2011; Sambasivan et al., 2011; von Maltzahn et al., 2013) and Ddx6 (p54/RCK), an RNA helicase found enriched in P-bodies and stress granules (Buchan and Parker, 2009). Pax7 and Ddx6 are expressed in QSCs and decrease during activation and commitment (Crist et al., 2012; Zammit et al., 2006). A density map of Pax7 and Ddx6 expression shows a broad range of expression levels across the QSC pool (Figure 1A-1C). A bivariate plot shows a positive correlation between Pax7 and Ddx6 (Figure S1A). Therefore, based on the expression profile of two different markers, the QSC pool is composed of a continuum of molecular states at tissue homeostasis. This supports models inferred from scRNA-seq analysis on SCs (De Micheli et al., 2020; Dell’Orso et al., 2019; Kimmel et al., 2020).

**Figure 1.**
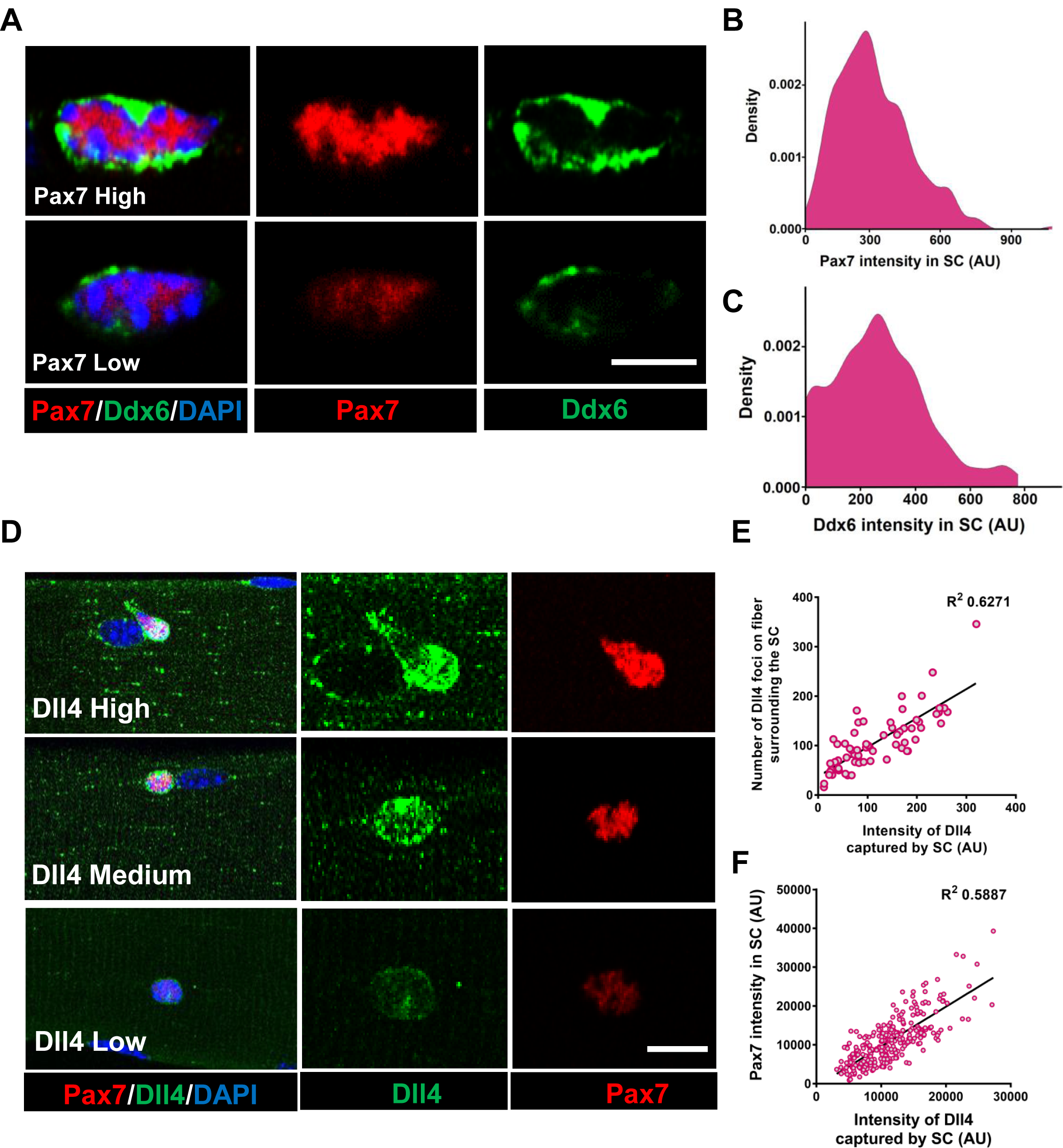
Spatial heterogeneous expression of Dll4 on the fiber correlates with SC diversity. (A) Representative images of a high and low Pax7^+^ SC that has corresponding high and low levels of Ddx6 protein. (B) A density map of Pax7 intensity in QSCs (n=3). (C) A density map of Ddx6 intensity in QSCs (n=3). (D) Representative images of regions of muscle fiber expressing heterogeneous levels of Dll4 protein and the corresponding expression of Pax7 in the adjacent SC. (E) A bivariate plot between the number of Dll4 foci on fiber adjacent to a SC and the intensity of Dll4 captured by the stem cell (n>3 mice). (F) An XY graph on isolated wildtype QSCs shows positive correlation between intensity of Dll4 captured by SC and its Pax7 intensity (n=3 mice). Scale bars, 5μm in (A) and 10μm in (D).

To determine the fate of cells along the cell state continuum, we isolated Pax7^high^, Pax7^medium^ and Pax7^low^ -expressing populations of SCs from Pax7-nGFP reporter mice (Rocheteau et al., 2012), and analyzed their potential for cell cycle entry and differentiation. As expected, the Pax7^low^ population entered cell cycle and differentiated faster than the Pax7^high^ expressers, suggesting that Pax7^high^ SCs are in a more dormant state and the Pax7^low^ SCs exist in a primed state (Rocheteau et al., 2012). The Pax7^medium^ population is in an intermediate molecular and phenotypic state between the Pax7^high^ and Pax7^low^ QSCs (Figure S1B-S1G). Therefore, the QSC pool exists as a continuum of phenotypic cell states.

### Heterogeneous expression of Dll4 in a multinucleated niche cell is coupled to QSC diversity

The multinucleated muscle fiber functions as a niche cell for the SC, regulating the depth of quiescence and rate of activation in response to injury (Eliazer et al., 2019). The Delta-Notch signaling pathway is an evolutionarily conserved inter-cellular signaling pathway for cell fate specification (Kopan and Ilagan, 2009). Adult muscle QSCs display active notch signaling (Bjornson et al., 2012; Fukada et al., 2011; Low et al., 2018; Mourikis et al., 2012; Verma et al., 2018). Overexpression of NICD1 increases Pax7 expression (Wen et al., 2012) and RBP-Jκ, a transcriptional co-activator of the notch signaling pathway, maintains the SC pool by repressing differentiation (Bjornson et al., 2012; Mourikis et al., 2012). It is not known if Notch ligands maintain the continuum of diverse cell states across the QSC pool.

We first examined Notch ligands on isolated single muscle fibers from postnatal day3 (p3), postnatal day7 (p7) and adult through microarray analysis. *Dll4* transcripts on the fiber showed the most enrichment as the SC transitioned from proliferating myogenic progenitors (postnatal) to becoming quiescent (adult) (Figure S2A). RT-qPCR on isolated adult single muscle fibers show the expression of *Dll4* transcripts, which was confirmed by RNAscope and immunofluorescence (Figure S2B, S2C and Figure 1D) (Kann and Krauss, 2019). Dll4 transcript is expressed in less than 1% of QSCs (De Micheli et al., 2020; Kimmel et al., 2020), suggesting that the muscle fiber is a source of SC-bound Dll4. Since the QSC pool is distributed as a continuum of cell states, we hypothesized that Dll4 expression across the multi-nucleated niche cell is heterogeneous. Single muscle fibers stained with anti-Dll4 shows that Dll4 protein formed clusters along the muscle fiber and were enriched around the QSCs. Further analysis revealed heterogeneous spatial expression of Dll4 protein along the muscle fiber that correlated with the amount of Dll4 captured by the SCs. We observed a positive correlation between the amount of Dll4 foci on the fibers and the intensity of Dll4 present on the adjacent QSC (Figure 1D, 1E). In contrast, Dll4 transcript expression was uniform along the fiber (Figure S2C).

To directly link DLL4 localization with SC diversity, we isolated and stained SCs for Dll4 and Pax7. A bivariate plot of Pax7 and Dll4 expression within SCs, reveals a positive correlation between the Dll4 captured by the stem cell and Pax7 intensity (Figure 1F). Comparison between Dll4 protein and Pax7 levels using a Pax7-nGFP reporter, confirmed the positive correlation between Pax7 and Dll4 protein expression (Figure S1E, S1H). Quantification of Dll4 expression levels on fiber adjacent to SCs per single muscle fiber of an Extensor Digitorum Longus (EDL) muscle shows that less than 10% of fibers are exclusively Dll4^High^ or DLL4^low^, suggesting SC diversity is coordinated in a fiber-autonomous manner (Figure S1I). Therefore, spatial heterogeneity of niche-derived DLL4 is coupled with a continuum of diverse molecular and phenotypic states across the QSC population.

### Niche-derived Dll4 maintains SC diversity

To determine if Dll4 from the muscle fiber controlled SC diversity, we deleted *Dll4* (Hozumi et al., 2008) specifically in the adult muscle fibers using a tamoxifen (tmx)-inducible HSA^CreMER^ mouse line (McCarthy et al., 2012) (Transgenic mice are herein called as MF-Dll4^fl/fl^) (Figure 2A). Dll4 ligand expression in the muscle fiber and the ligand captured by the SCs decreased in MF-Dll4^fl/fl^ compared to control, suggesting that the muscle fiber is the major source of SC-bound Dll4 in uninjured muscle (Figure 2B-2D). It is possible that cell sources other than the muscle fiber could provide Dll4 to SCs in other contexts (Verma et al., 2018). We observed that Pax7 and Ddx6 expression in SCs from MF-Dll4^fl/fl^ fibers decreased compared to controls (Figure 2E, 2G). Density maps show that there is a shift in the distribution of expression levels of Pax7 and Ddx6. We calculated the variance of the expression levels to gauge the spread of the distribution. The decrease in variance in MF-Dll4^fl/fl^ compared to controls, indicates reduced diversity across the QSC pool, effectively reducing the range of states on the continuum (Figure 2F, 2H). We also observed a 50% decrease in the total number of Pax7^+^ SCs in MF-Dll4^fl/fl^ fibers compared to controls (Figure 2I), suggesting a stem cell loss phenotype. A decrease in SC diversity, number and Pax7 levels can be explained by a loss of Pax7^high^ SCs (due to apoptosis or fusion), or a shift in the continuum towards a more committed fusion-competent state, followed by fusion of SCs expressing the lowest levels of Pax7. qRT-PCR on QSCs from control and MF-Dll4^fl/fl^ reveals a decrease in *pax7* (stem cell marker) and an increase in *myoD* (activation marker) and *myogenin* (differentiation marker) transcripts (Figure 3B). The simultaneous loss of Pax7 and induction of myogenin, a gene not normally expressed in QSCs, suggests a shift in the QSC continuum. While a loss of Pax7^high^ cells alone would not increase Myogenin expression, we cannot exclude a loss of Pax7^high^ SCs in addition to the shift in continuum. Deletion of Dll4 in Pax7^+^ SCs using a Pax7^CreER^ transgenic mouse line (Nishijo et al., 2009) (Transgenic mice are referred to as SC-Dll4^fl/fl^) did not change Pax7^+^ SC number, compared to controls (Figure S3A, S3B), suggesting that Dll4 does not have an autocrine role in QSCs. Together, these results suggest that Dll4 from the muscle fiber is required to maintain the continuum of diverse cellular states across the QSC pool.

**Figure 2.**
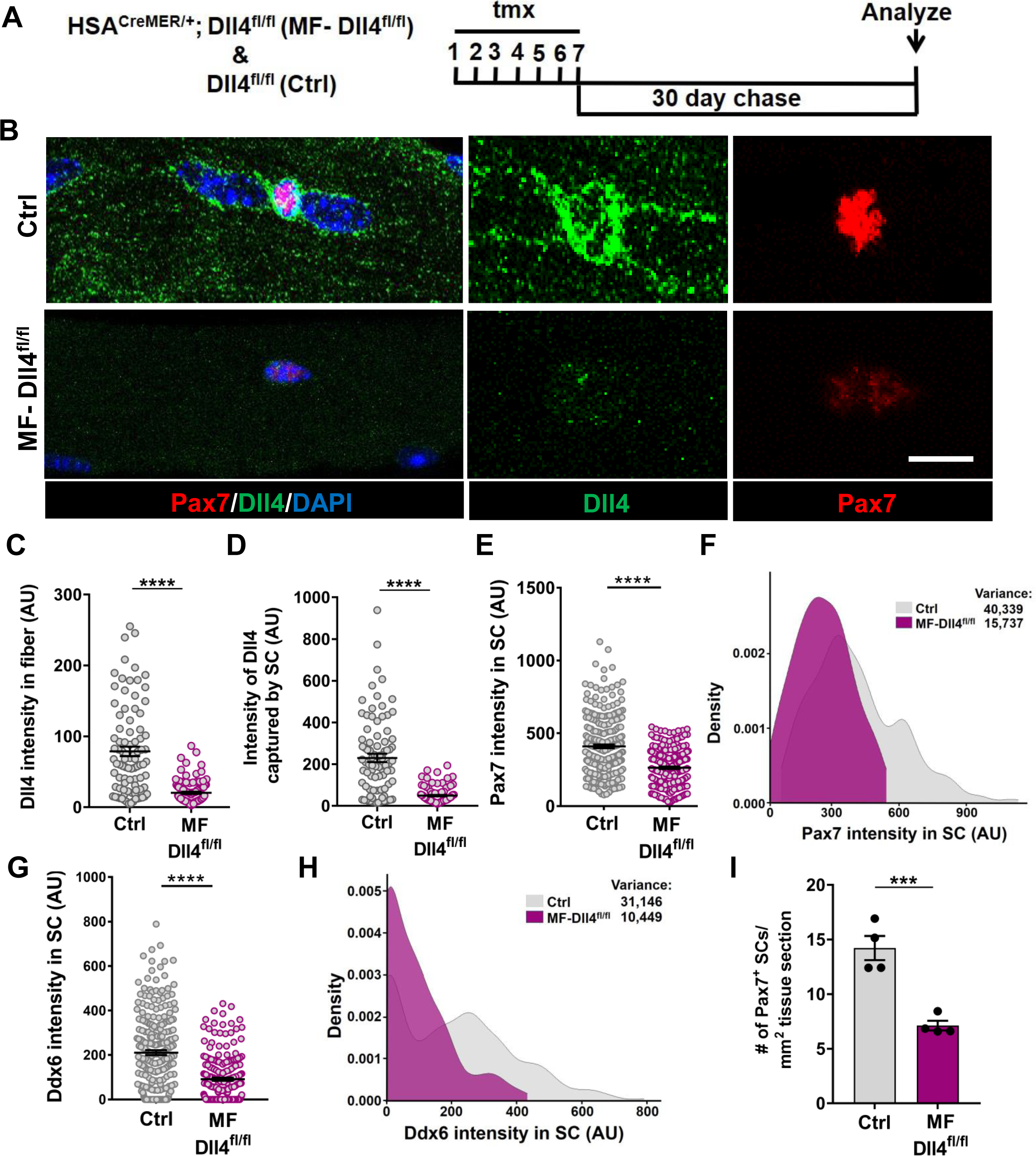
Muscle fiber derived Dll4 maintains a continuum of diverse states in the QSC pool. (A) Schematic representation of the experimental design. (B-F) Representative images of Dll4 and Pax7 expression in Control and MF-Dll4^fl/fl^ fibers and SCs respectively (in B) and quantification of Dll4 intensity in fibers (in C), Intensity of Dll4 captured by SCs (in D), Pax7 intensity in SCs (in E) and a density map of Pax7 intensity in SCs (in F) (n=3 mice). (G, H) Ddx6 intensity in SCs (in G) and density map of Ddx6 intensity in SCs (in H) on Control and MF-Dll4^fl/fl^ fibers (n=3 mice). (I) Number of Pax7^+^ SCs in tissue sections of Control and MF-Dll4^fl/fl^ fibers (n=4 mice). Error bars, mean ± s.e.m; ***p<0.001, ****p<0.0001; Scalebars, 10 μm in (B).

**Figure 3.**
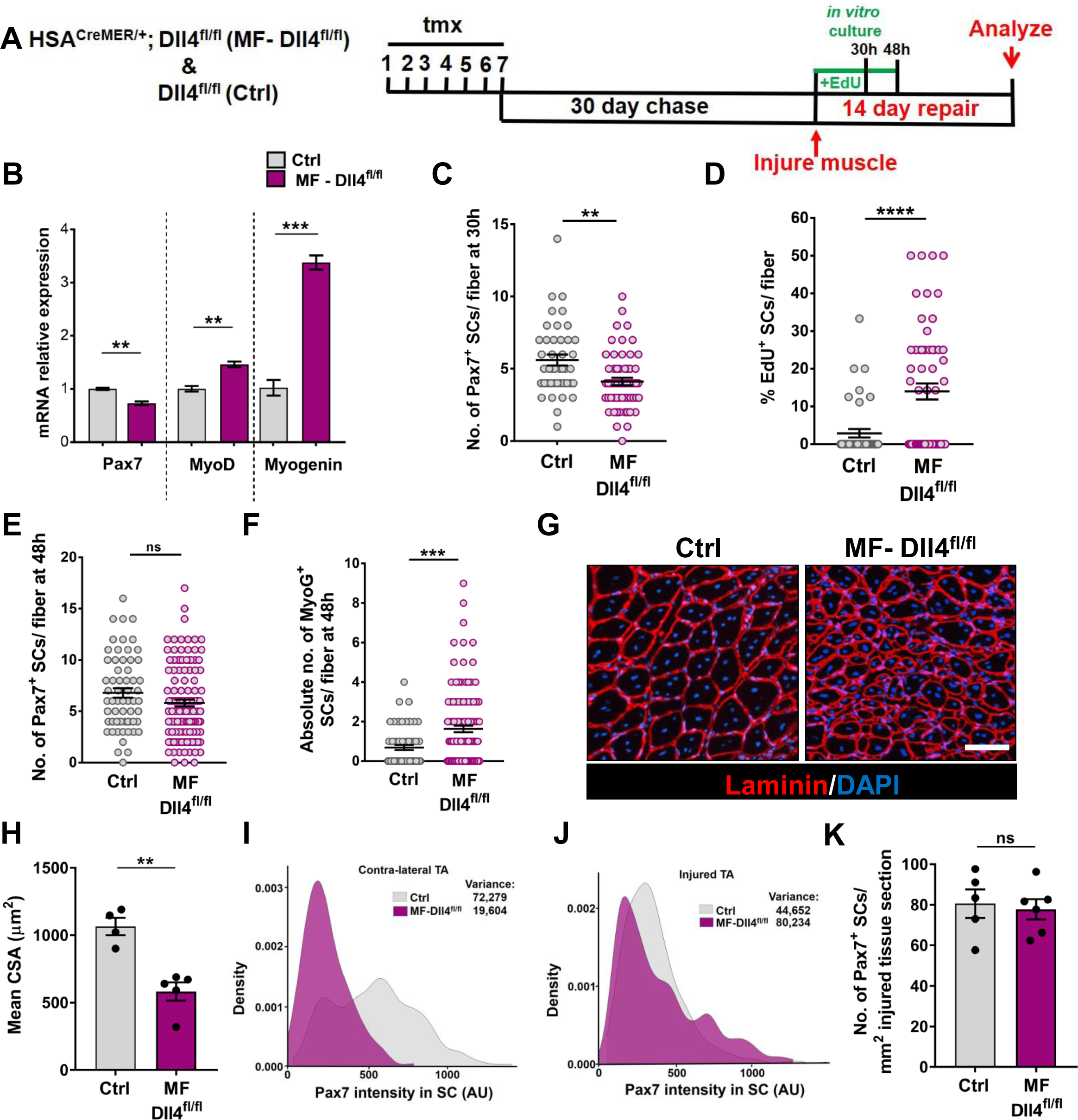
Niche derived Dll4 constrains the proliferative and commitment potential of the QSC pool. (A) Schematic representation of the experimental design. (B) qRT-PCR for *Pax7, MyoD* and *Myogenin* transcripts in freshly isolated SCs from Control and MF-Dll4^fl/fl^ fibers (n=3). (C, D) Control and Dll4 deleted fibers were cultured in plating media containing EdU for 30h, the number of Pax7^+^ SCs (in C) and percent of EdU^+^ SCs (in D) per fiber were quantified (n=3). (E, F) Control and MF-Dll4^fl/fl^ fibers were cultured *in vitro* for 48h, the number of Pax7^+^ SCs (in E) and the MyoG^+^ cells (in F) per fiber were quantified (n=3). (G, H) Representative images (in G) and quantification (in H) of mean cross-sectional area of Control and MF-Dll4^fl/fl^ TA fibers, injured and regenerated for 14 days (n≥4). (I, J) Density map of Pax7 expression in SCs from Ctrl and MF-Dll4^fl/fl^ contra-lateral TA (in I) and injured TA (in J), 14d post injury (n=3). (K) Number of Pax7^+^ SCs in regenerated TA muscle, 14d after injury (n≥5). Error bars, mean ± s.e.m; **p<0.01, ***p<0.001, ****p<0.0001; Scale bars, 100μm in (G).

### Dll4 from the muscle fiber constrains the proliferative and commitment potential across the QSC pool

To analyze the phenotypic states across the QSC pool that remains on the Dll4-depleted niche, we cultured isolated single muscle fibers from control and MF-Dll4^fl/fl^ mice for 30h (Figure 3A). SCs on a Dll4-depleted niche entered cell cycle faster than controls (Figure 3C, 3D). This is consistent with a loss of more dormant Pax7^high^ SCs. We next cultured the SCs on control and Dll4 depleted niche for 48h in 10% serum-containing media (Figure 3A). The number of SCs on a Dll4-depleted niche was similar to controls, suggesting compensatory proliferation of the remaining SCs (Figure 3E). The absolute number of Myogenin^+^ cells per fiber was increased, suggesting that a fraction of the remaining SCs on a Dll4-depleted niche rapidly entered the differentiation program when exposed to mitogen. (Figure 3F). Therefore, in the absence of fiber-derived Dll4 the QSC pool shifts along a continuum, losing its dormant features and becoming more proliferative and committed. Induction of *myogenin* is consistent with the anti-differentiative role of RBP-Jκ (Bjornson et al., 2012; Mourikis et al., 2012). Based on prior work, we would not have predicted an increase in SC proliferation in mitogen after niche depletion of Dll4 (Bjornson et al., 2012; Mourikis et al., 2012).

We next examined the effect of a Dll4-depleted niche on fate potential of the remaining SCs in response to muscle injury. Using the tmx-inducible HSA^CreMER/+^ genetic model, the niche is genetically modified, but the SCs are genetically wildtype. The wildtype SCs reform the adult muscle fibers during regeneration (Eliazer et al., 2019). Therefore, any regenerative phenotype observed is directly due to the loss of Dll4 in the niche prior to injury. Fourteen days after injury, muscle fiber size in MF-Dll4^fl/fl^ TA muscle was significantly smaller than controls (Figure 3G, 3H), consistent with the rapid entrance of some SCs into the differentiation program, as reported after deletion of RBP-Jκ in SCs (Bjornson et al., 2012; Mourikis et al., 2012). The inability to repair muscle fibers after a loss in SC diversity suggests a reduced pool of fusion-competent progenitors. The shift in the continuum after Dll4 deletion, suggests a role for Pax7^high^ SCs in the generation of fusion-competent progenitors.

We next asked whether the remaining population of Pax7^+^ SCs in a Dll4-depleted niche was capable of reestablishing the normal continuum of Pax7^+^ states in the injury model. In contrast to the contra-lateral uninjured muscle (Figure 3I), the average Pax7 levels and the variance across the SC population was not different in a Dll4-replenished niche and control niche (Figure 3J). In addition, the number of Pax7^+^ cells were similar to control regenerated muscle (Figure 3K). This was unanticipated due to the decrease in Pax7 levels and a shift along the continuum towards a more committed state in the QSC pool (Kuang et al., 2007; Rocheteau et al., 2012). Therefore, SCs expressing modest levels of Pax7 have the potential to produce SCs with higher levels of Pax7 during muscle regeneration, revealing an unappreciated level of plasticity and bidirectionality along the SC continuum.

### Mib1 directs spatial patterning and activation of Dll4

Single nucleus RNA sequencing has uncovered substantial transcriptional diversity across muscle fibers (Kim et al., 2020; Petrany et al., 2020). However, Dll4 activity is regulated post-translationally. Mindbomb1 (Mib1), is an E3 ubiquitin ligase that activates all Notch ligands in the signal sending cell by adding mono-ubiquitin groups onto the ligands (Koo et al., 2005; Koo et al., 2007). The hypothalamus-pituitary-gonadal axis has been shown to induce Mib1 in muscle fibers during development, thus activating notch signaling in juvenile SCs (Kim et al., 2016). To determine if the heterogeneous pattern of Dll4 on the adult muscle fiber is directed by Mib1, we analyzed the spatial expression of Mib1 on adult muscle fibers. We stained isolated single muscle fibers with Mib1 antibody and found heterogeneous expression pattern of Mib1 on individual muscle fibers. Similar to Dll4 expression, we observed a positive correlation between the intensity of Mib1 expression on muscle fibers and the Pax7 intensity within the adjacent SC (Figure 4A, 4B). Formation of Dll4 clusters is a marker of the activated form of the ligand (Le Borgne and Schweisguth, 2003). To test whether Mib1 played a role in clustering and activating Dll4 on the muscle fibers, Mib1 floxed mice (Koo et al., 2007) were crossed with tamoxifen inducible HSA^CreMER^ mice (Tg: hereafter called as MF-Mib1^fl/fl^). After a 30-day chase, the expression of *Mib1* transcripts were decreased in the Mib1-depleted myofibers compared to controls (Figure S4A, S4B). Analysis of Dll4 expression in control and MF-Mib1^fl/fl^ isolated single muscle fibers, reveals a loss of Dll4 clusters and a reduction in the intensity of Dll4 expression in Mib1-depleted niche (Figure 4C-4E). Therefore, Mib1 directs the activation and spatial patterning of Dll4 within a multinucleated niche cell.

**Figure 4.**
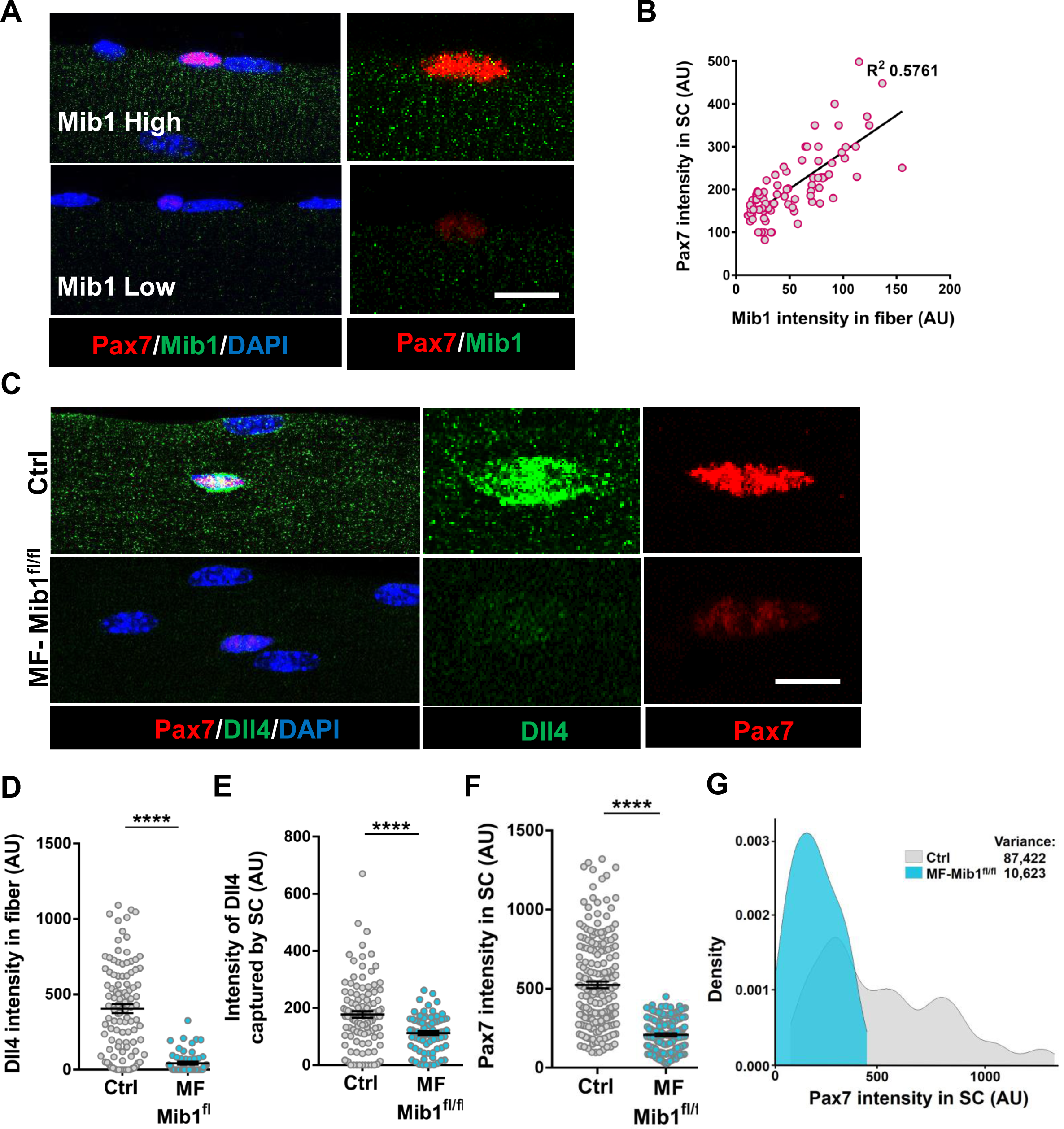
Niche-derived Mib1 directs spatial patterning and activation of Dll4. (A) Representative images of WT fibers showing regions of high and low Mib1 expression and the adjacent Pax7^+^ SC. (B) A bivariate plot showing Mib1 intensity in the fiber and Pax7 intensity in the adjacent SC. (C-G) Representative images of Dll4 expression in fibers and Pax7 expression in SCs present on Control and MF-Mib1^fl/fl^ fibers (in C), quantification of Dll4 intensity in fibers (in D), Intensity of Dll4 captured by SCs (in E), Pax7 intensity in SCs (in F) and a density map of Pax7 intensity in SCs (in G) on Control and MF-Mib1^fl/fl^ fibers (n=3 mice). Error bars, mean ± s.e.m; ****p<0.0001; Scale bars, 5μm in (A) and (C).

### Niche-derived Mib1 maintains a continuum of states across the QSC pool

Based on the loss of SC-bound Dll4 after Mib1 deletion in the niche, we analyzed Pax7 levels and diversity across the QSC pool on a Mib1-depleted niche. Compared to controls, Pax7 levels decreased, the variance across the QSC pool was reduced and SC number was less in a Mib1-depleted niche (Figure 4F, 4G, Figure S4C). RT-qPCR of myogenic markers show that *Pax7* and *myoD* transcript levels did not change. However, *myogenin* was upregulated in the SCs remaining on a Mib1-depleted niche (Figure S4D).

In response to activation cues, SCs on a Mib1-depleted niche proliferate faster and upregulate Myogenin compared to the SCs on control muscle fibers, indicating that the remaining SCs are primed for proliferation and differentiation (Figure S4E-S4G). In response to muscle injury, the size of regenerating muscle fibers was smaller than the control muscle fibers (Figure S4H, S4I). Therefore, niche-derived Mib1 expressed in a spatially heterogenous pattern in a fiber-autonomous manner maintains the normal continuum of diverse QSC states during tissue homeostasis.

## Discussion

The multinucleated muscle fiber acts as a niche cell to maintain the QSC pool in a continuum of diverse states through the spatial patterning of the E3 ligase, Mib1 and activation of Dll4, in a fiber-autonomous manner. The composition and location of niche cells are critical for stem cell regulation. Our study reveals that the localization of niche factors within multinucleated cells are an important component of stem cell regulation.

The QSC pool is heterogeneous and exists in an array of cellular states based on SC markers. It is important to actively maintain the diversity of cell states so that in the event of an injury, all the SCs are not equally lost to activation or differentiation. We demonstrate the existence of a continuum of quiescent cell states ranging from deep quiescent, non-committed states to more committed states. This continuum is actively maintained in uninjured muscle by the spatial pattern of Dll4 and Mib1 on fiber-autonomous levels. Depletion of Dll4 and Mib1 in muscle fibers causes a contraction of the SC continuum towards more homogeneous primed states.

The spatial localization of the different factors in the cytoplasm of a single multinucleated cell governs the cell state of the QSC. We believe that the expression of Wnt4 along the fiber is ubiquitously expressed, thereby inhibiting proliferation of all QSCs. The heterogenous spatial expression of Dll4 within the muscle fiber allows the maintenance of a continuum of differentiated states, with SCs in a region of low Dll4 expression primed and ready to differentiate when subjected to injury *in vivo* or exposed to mitogen *in vitro*.

Although snRNA sequencing shows transcriptional heterogeneity within the myonuclei of single muscle fibers (Kim et al., 2020; Petrany et al., 2020), we show that the transcripts of Dll4 are uniformly expressed along the muscle fiber and the protein expression is heterogeneous. This highlights the importance of protein heterogeneity within single muscle fibers. The varied expression of Dll4 is established by the E3 ligase, Mib1 that adds ubiquitin groups to Dll4 and activates them by a process of endocytosis (Koo et al., 2005; Koo et al., 2007). The formation of endosomes might play a role in clustering the Dll4 ligand to present them to the adjacent receptors on the QSCs.

The deletion of Mib1 or Dll4 within the fiber resulted in a loss of 50% of the SC pool, whereas, deletion of RBP-Jκ in adult SCs displayed a more robust depletion of the SC pool (Bjornson et al., 2012; Mourikis et al., 2012). This suggests that RBP-Jκ might have targets other than the Notch signaling pathway. Deletion of RBP-Jκ in SCs also caused the SC pool to precociously differentiate. There was some evidence of proliferation, but not in those SCs that eventually differentiated, suggesting a divergent response in some SCs compared to others. Why did deletion of RBP-Jκ within the QSC pool result in two different phenotypes? We propose that SCs sit along a continuum dictated by the levels of fiber-derived Dll4 captured on the SC. Disruption of Dll4 and Mib1 in the muscle fiber re-equilibrates where each SC sits on that continuum. With the response dependent on the relative position along the continuum.

scRNA-seq on cells from regenerating muscle suggests that Dll1 is expressed in a subset of differentiating, Myogenin^+^ cells and is important for self-renewal (Yartseva et al., 2020). *In vitro* experiments with co-culture of primary myoblasts with 3T3 cells overexpressing Dll4 (Low et al., 2018), or with endothelial cells (Verma et al., 2018) suggests the importance of the Notch ligand Dll4 in self-renewal. Our results provide the first direct demonstration of a Notch ligand from a specific cell type that is critical to maintain the adult SC pool in a quiescent and non-differentiated state in vivo. Notch ligands added to SCs in vitro promoted quiescence of myoblasts, but did not restore stemness (Sakai et al., 2017). Therefore, quiescence is not equivalent to stemness; consistent with the observations that only subsets of QSCs possess self-renewal potential in transplantation assay (Chakkalakal et al., 2012; Rocheteau et al., 2012; Sacco et al., 2008).

Prior work demonstrated that SCs with low levels of Pax7 are less competent for self-renewal in vivo. Based on the decrease in Pax7 levels after Dll4 depletion from the niche, we would have predicted a diminution of the SC pool in response to injury. Instead Pax7 levels were increased in the SC pool after injury. Therefore, in the context of muscle regeneration, there is a level of plasticity in the SC pool. We do not favor the alternative interpretation that another source of cells replenishes the Pax7^+^ SC compartment.

In the future, it would be interesting to investigate spatial heterogeneity of Notch ligands in other multinucleated cells and other stem cell niches. Our findings provide proof-of-principle for niche-based strategies to modify the fate of stem cells while residing in a quiescent state.

---

**Figure S1.**
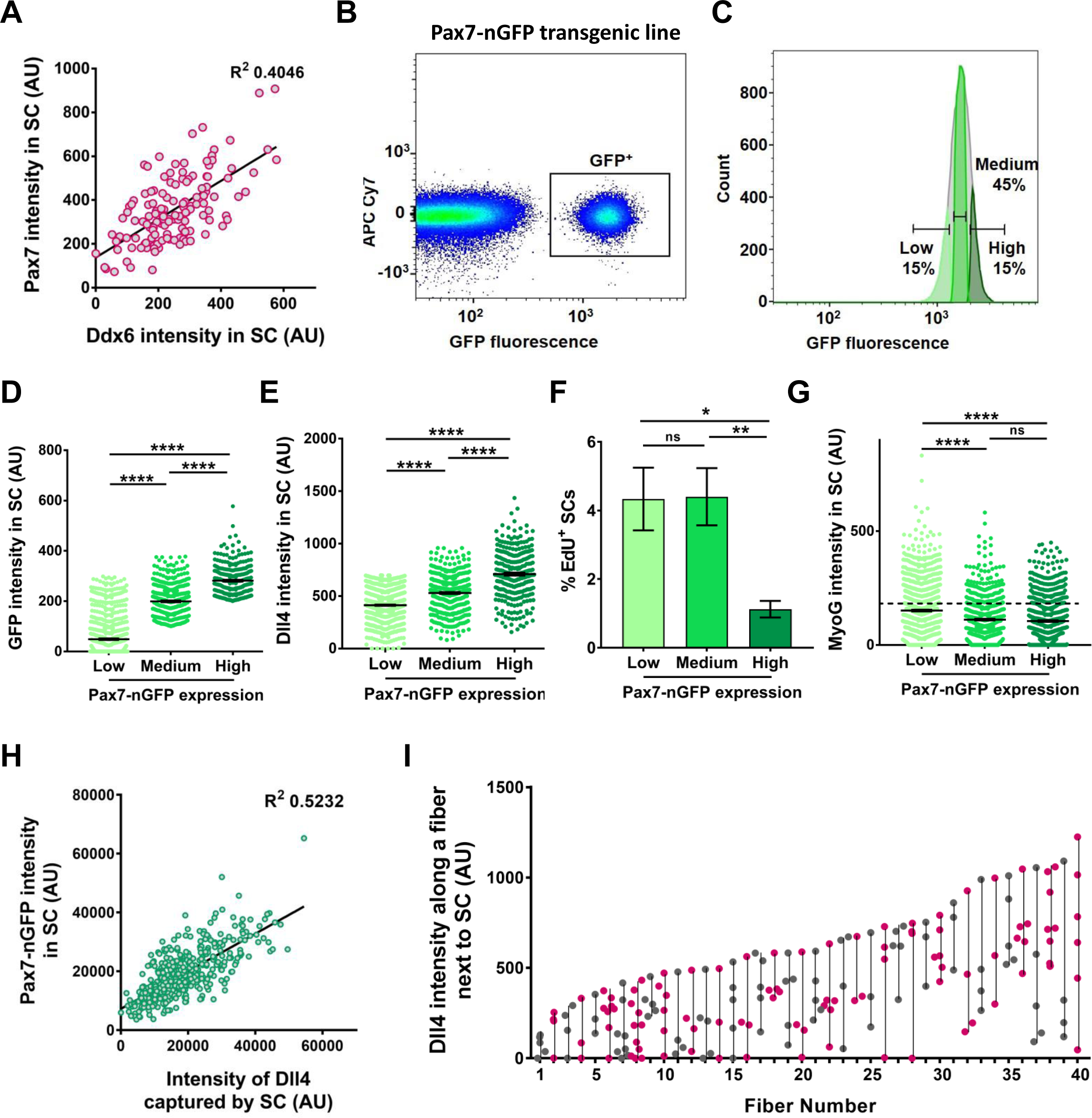
Adult quiescent SCs exist in a continuum of diverse molecular and phenotypic states. (A) A bivariate plot between Pax7 and Ddx6 intensity in QSCs (n=2). (B, C) FACS sorting strategy of obtaining GFP+ SCs from a Pax7-nGFP transgenic reporter line (in B) and isolation of GFP low, medium and high expressing SCs (in C). (D-G) The three sorted populations were fixed immediately and stained for GFP (in D), Dll4 (in E), *in vitro* cultured and treated with EdU for 42 hours (in F) and cultured *in vitro* in low serum for 3 days and stained for Myogenin protein (in G) (n=3). (H) An XY plot of freshly isolated SCs from Pax7-nGFP transgenic mouse line shows positive correlation between intensity of Dll4 captured by SC and its GFP (Pax7) intensity (n=3 mice). (I) Dll4 expression along muscle fibers. Each vertical line represents a single fiber and each dot on the line represents the Dll4 intensity in the fiber adjacent the SC. Error bars, mean ± s.e.m; *p<0.05, **p<0.01, ****p<0.0001.

**Figure S2.**
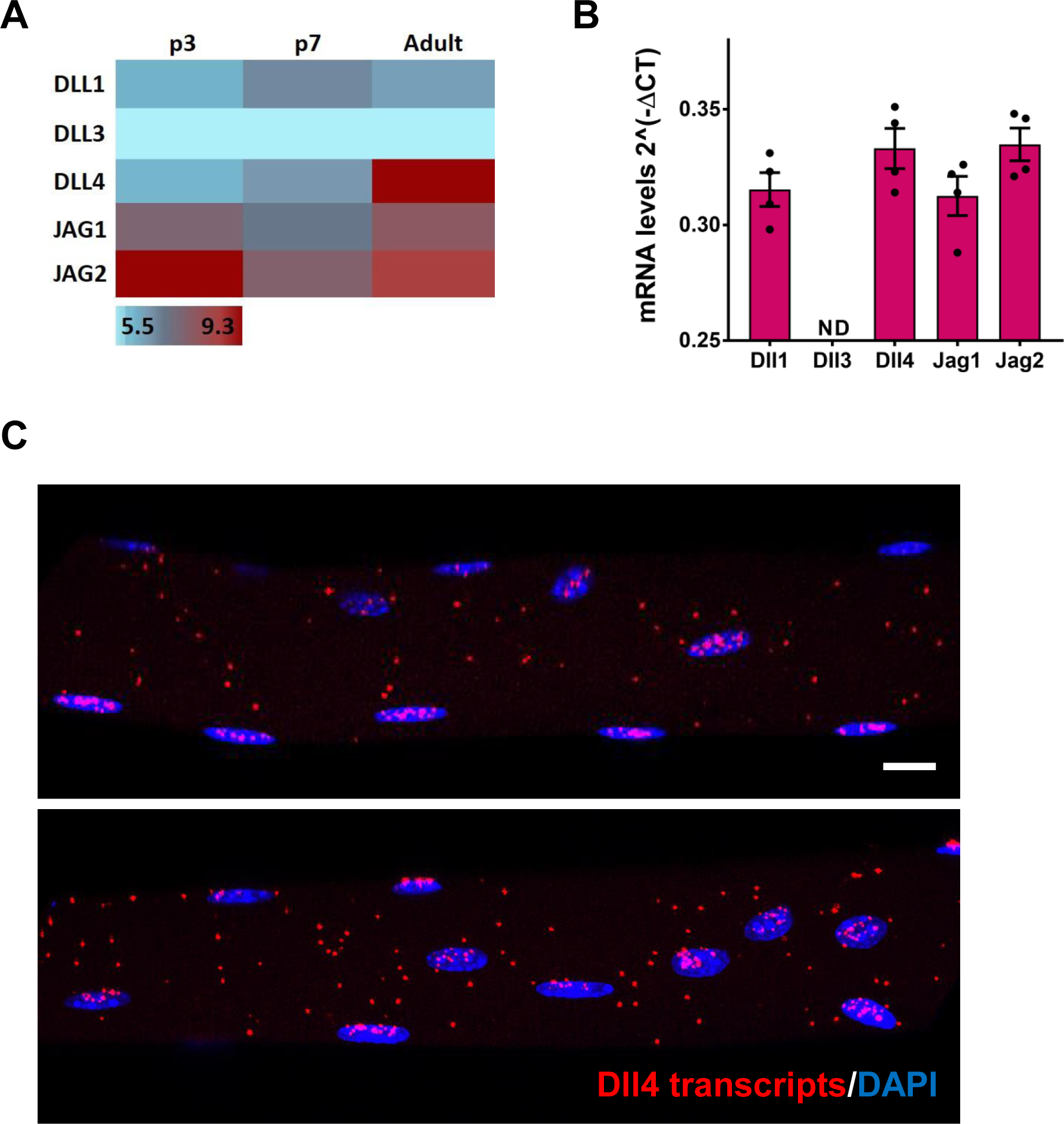
Dll4 ligand is highly expressed in adult muscle fibers. (A) Heatmap of Notch ligands from microarray of muscle fibers isolated from postnatal day 3 (p3), day 7 (p7), and adult Extensor Digitorum Longus (EDL) muscle (n=2), The expression values are log2 transformed and range from 5.5 (low, blue) to 9.3 (high, red). (B) qRT-PCR of Notch ligands in isolated adult muscle fibers. The notch ligand transcripts are normalized to GAPDH (n=4); ND, not detected. (C) RNAscope of Dll4 transcripts on single adult muscle fibers (n=3). Error bars, mean ± s.e.m; Scale bars, 10μm in (C).

**Figure S3.**
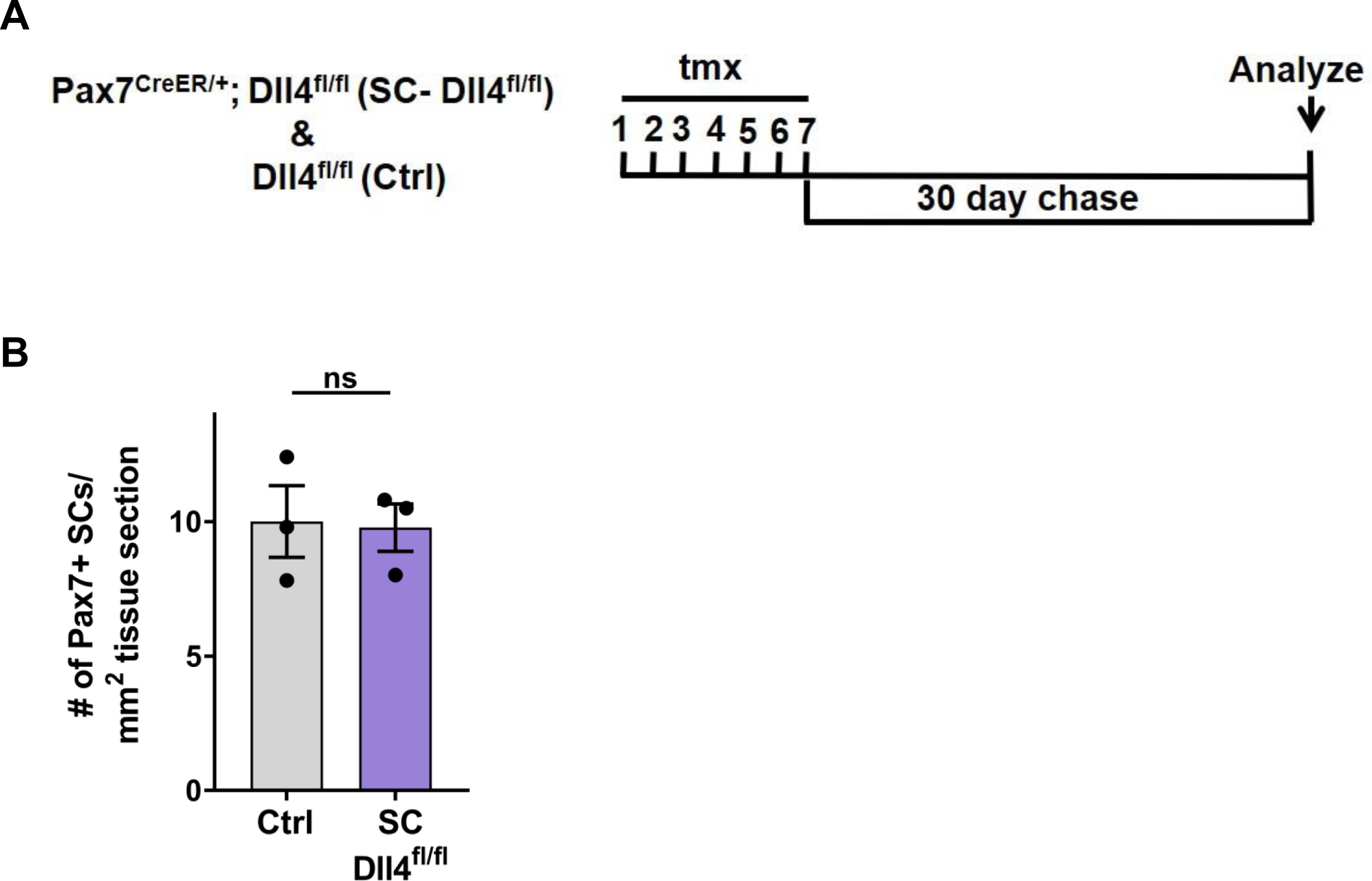
Dll4 does not have a cell autonomous role in QSC. (A) Schematic representation of the experimental design. (B) Number of Pax7^+^ SCs in TA muscle sections of Control and SC-Dll4^fl/fl^ fibers (n=3 mice). Error bars, mean ± s.e.m; ns, non-significant.

**Figure S4.**
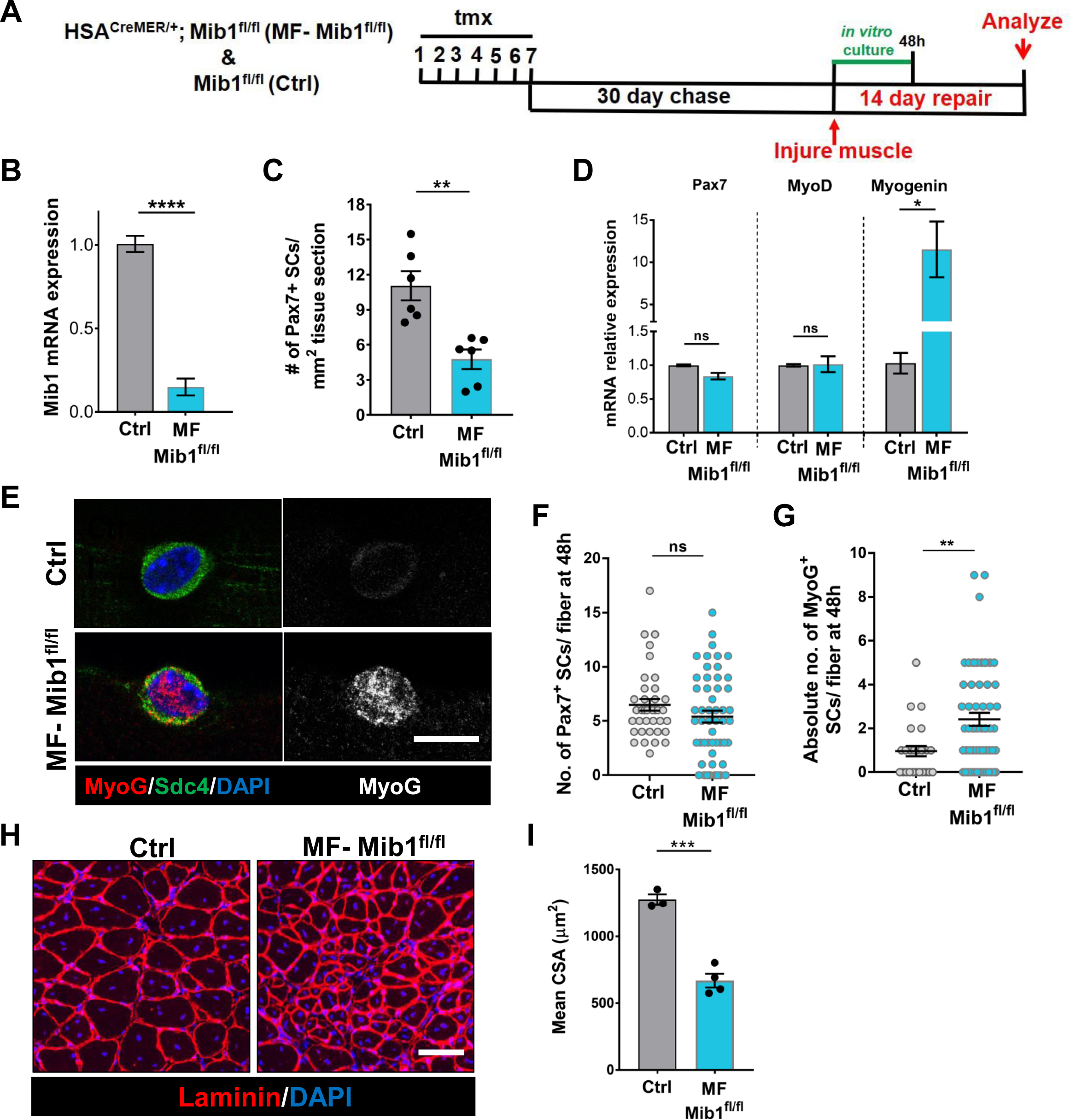
Loss of niche-derived Mib1 phenocopies Dll4 mutant phenotype. (A) Schematic representation of the experimental design. (B) qRT-PCR for *mib1* transcripts in fibers of Control and MF-Mib1^fl/fl^ normalized to GAPDH (n=3). (C) Number of Pax7^+^ SCs in TA muscle sections of Control and MF-Mib1^fl/fl^ (n=6). (D) qRT-PCR for *Pax7, MyoD* and *Myogenin* transcripts in freshly isolated SCs from Control and MF-Mib1^fl/fl^ fibers, normalized to GAPDH (n=3). (E-G) Representative images of Sdc4 and MyoG expression in SCs from Control and Mib1 deleted fibers, cultured *in vitro* in plating media for 42h (in E), the number of Pax7^+^ SCs (in F) and the number of MyoG^+^ cells (in G) per fiber were quantified (n=3). (H, I) Representative images (in H) and quantification (in I) of mean cross-sectional area of Control and MF-Mib1^fl/fl^ TA muscle fibers, injured and regenerated for 14 days (n≥<u>3</u>). Error bars, mean ± s.e.m; *p<0.05, **p<0.01, ***p<0.001, ****p<0.0001; Scale bars, 10 μm in (E) and 100μm in (H).

**Table S1.** Related to Figure 1. Microarray expression of Notch ligands in p3, p7 and Adult muscle fibers (log2 values)

**Microarray expression of Notch ligands in p3, p7 and Adult muscle fibers (log2 values)**
|  | **p3** | **p7** | **Adult** |
| --- | --- | --- | --- |
| **Dll1** | 7.179268 | 7.864147 | 7.530619 |
| **Dll3** | 5.568594 | 5.568594 | 5.568594 |
| **Dll4** | 7.124728 | 7.676524 | 9.269889 |
| **Jag1** | 8.456506 | 8.075812 | 8.670788 |
| **Jag2** | 9.312074 | 8.530521 | 9.012903 |

**Table S2.**
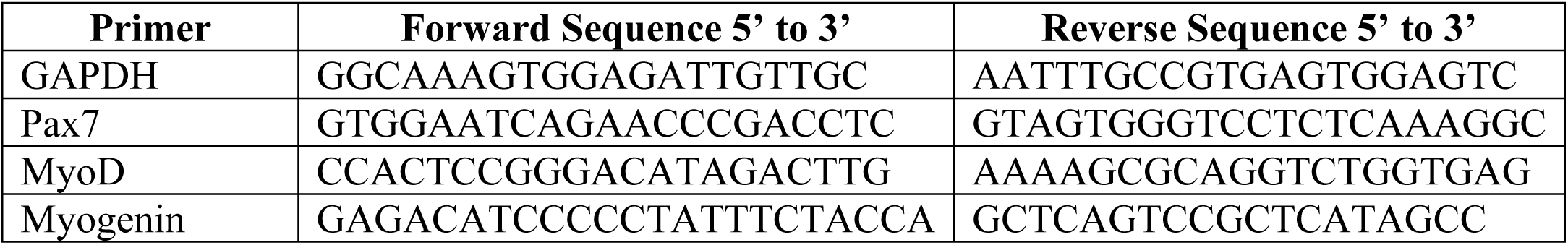
Related to STAR Methods. Primers used for qRT-PCR analysis

## Acknowledgements

We would like to thank Drs. Karyn Esser, Charles Keller, Young-Yun Kong and Sonoko Habu for providing transgenic mice. We would like to thank Hallie Nelson for technical assistance, and members of the Brack laboratory for critical discussions during the preparation of this manuscript. We acknowledge the UCSF Parnassus Flow Cytometry Core (RRID:SCR_018206) supported in part by Grant NIH P30 DK063720 and by the NIH S10 Instrumentation Grant S10 1S10OD021822-01. This work was supported by NIH grants (R01AR060868, R01AR061002, R01AR076252) to A.S.B., and (F32AR067594) to S.E.

## Author contributions

S.E. conceived the project, designed and performed experiments, analyzed data, interpreted results, and wrote the manuscript. X.S. performed experiments and analyzed data. A.S.B. conceived the project, designed and interpreted experiments, and edited the manuscript.

## STAR Methods

### LEAD CONTACT AND MATERIALS AVAILABILITY

Further information and requests for resources and reagents should be directed to and will be fulfilled by the Lead Contact, Andrew S. Brack.

### EXPERIMENTAL MODEL AND SUBJECT DETAILS

#### Animals

Mice were housed and maintained in accordance with the guidelines of the Laboratory Animal Research Center (LARC) of University of California, San Francisco. C57BL/6 were obtained from Jackson Laboratory. Previously published Pax7-nGFP (Rocheteau et al., 2012), HSA^CreMER^ (McCarthy et al., 2012), Dll4^flox/flox^ (Hozumi et al., 2008), Mib1^flox/flox^ (Koo et al., 2007), Pax7^CreER^ (Nishijo et al., 2009) were used in this study. All mice used for experiments were adults, between 12-16 weeks of age unless otherwise indicated. The control and experimental mice used are littermates in all experiments. Approximately equal numbers of male and female mice were used in all experiments. Animals were genotyped by PCR using tail DNA. Primer sequences are available upon request.

## METHOD DETAILS

### Animal Procedures

Tamoxifen (Sigma) was dissolved in corn oil at a concentration of 20mg/ml. Both control and experimental mice were administered tamoxifen at a concentration of 150mg/kg per day for seven continuous days by intraperitoneal injection. The mice were left to chase for 30 days before analysis.

### Muscle Injury

Control and experimental mice were anesthetized by isofluorane inhalation and 50µl of 1.2% BaCl_2_ was injected into and along the length of the tibialis anterior (TA) muscle. After 14 days of regeneration, mice were euthanized, the contra-lateral uninjured TA and injured TA muscle were fixed immediately in 4% PFA and frozen in 20% sucrose/OCT medium. 8µm cross-sections of the muscle were made and stained for anti-laminin. 10x images were collected at three regions in the mid-belly of each muscle. Only mice that had more than 80% injury in their TA were analyzed. All the regenerating fibers in the entire TA section were analyzed for fiber size. The average cross-sectional area of the fibers was determined using ImageJ software.

### Isolation of Single Muscle Fibers

Single muscle fibers were isolated from the Extensor Digitorum Longus (EDL) muscle of the adult mouse as described previously (Eliazer et al., 2019). The single fibers were fixed immediately in 4% PFA for 10 minutes or cultured in plating media (DMEM with 10% horse serum).

For *in vitro* cell cycle entry assays, single muscle fibers from control and Myofiber-Dll4^fl/fl^ mice were harvested and cultured in plating media containing EdU (10µm; Carbosynth) for 30 hours. For *in vitro* differentiation assay, the single muscle fibers from control, Myofiber-Dll4^fl/fl^ and Myofiber-Mib1^fl/fl^ were cultured in plating media for 48 hours. The fibers were then fixed and stained for different antibodies. EdU staining was done using the Click-iT Plus EdU Alexa Fluor™ 594 Imaging Kit (Invitrogen) followed by staining with anti-Pax7 antibody.

### Isolation of SCs and Fluorescence Assisted Cell Sorting (FACS)

Satellite cells were isolated from hindlimb and forelimb muscles as previously described(Eliazer et al., 2019). The mononuclear muscle cells were stained for PE-Cy7 anti-mouse CD31 (clone 390; BD Biosciences), PE-Cy7 anti-mouse CD45 (clone 30-F11; BD Biosciences), APC-Cy7 anti-mouse Sca1 (clone D7; BD Biosciences), PE anti-mouse CD106/VCAM-1 (Invitrogen) and APC anti-α7 integrin (clone R2F2; AbLab). FACS was performed using FACS Aria II (BD Biosciences) by gating for CD31^-^/CD45^-^/Sca1^-^/α7 integrin^+^/VCAM1^+^ to isolate SCs. SCs from Pax7-nGFP mouse were sorted for GFP fluorescence. The GFP^+^ gate was divided into top 15% (GFP^high^ fraction), middle 45% (Pax7^medium^ fraction) and bottom 15% (Pax7^low^ fraction). The isolated SCs were fixed immediately at t0 or cultured in growth media (Ham’s F10 media, 20% FBS, 5ng/ml FGF2) containing 10μm EdU for 42h (cell cycle entry assay) or cultured in plating media for 3 days (differentiation assay).

### Immunostaining

Fixed myofibers were permeabilized with 0.2% TritonX-100/PBS and blocked with 10% goat serum/0.2% TritonX/PBS. Primary antibodies used in this study were: mouse anti-Pax7 (DSHB), rabbit anti-Laminin (Abcam), rabbit anti-Dll4 (Thermo Fisher Scientific), rabbit anti-Mib1 (Sigma), rabbit anti-Syndecan4 (Abcam), rabbit anti-DDX6 (Bethyl Laboratories Inc.), rabbit anti-Myogenin (Santa Cruz Biotechnology) and DAPI (Life Technologies). Primary antibodies were visualized with fluorochrome conjugated secondary antibodies (Invitrogen). The stained fibers were mounted in Fluoromount-G mounting medium (SouthernBiotech). For most of the staining, the images were taken using a 20x Plan Fluor objective of the Nikon Eclipse Ti microscope. Anti-Pax7 and anti-Ddx6 stained images were obtained with a 40x Plan Fluor objective of the same microscope. Dll4 and Mib1 stained images were taken with 40x oil objective of Leica DMi8 Confocal Microscope. 15μm z-stacks around the Pax7^+^ SC were taken and the sum projection of the images were obtained. The filter settings, gain and exposure values were kept constant between experiments. The intensity of expression was determined by manually drawing a region of Interest (ROI) on a Pax7 positive SC. This will give the mean pixel intensity of the ROI in all the channels. The ROI is copied onto another region where there is no Pax7 positive cell to calculate the background intensity. The background intensity is subtracted and the mean intensity is plotted with GraphPad Prism 7. Representative images for antibody staining were taken using Leica DMi8 Confocal Microscope.

### RNAscope

RNAscope for *dll4* was performed on fixed myofibers as previously described (Kann and Krauss, 2019). Briefly the single muscle fibers are isolated, fixed with 4% PFA and dehydrated with 100% methanol. The fibers are then rehydrated with a series of decreasing concentrations of methanol and PBS with 0.1% Tween 20 followed by protease digestion, hybridization of the RNA probe, amplification of signal and conjugation with fluorophore.

### Quantitative PCR (qPCR)

Total RNA was isolated from single muscle fibers (from EDL muscle), or SCs using Trizol (Invitrogen) according to the manufacturer’s protocol. The RNA was DNAse treated using Turbo DNA free kit (Life Technologies). cDNA was synthesized from RNA using the Superscript First Strand Synthesis System (Invitrogen). qPCR was performed in triplicates from 5ng of RNA per reaction using Platinum SYBR Green qPCR Super Mix-UDG w/ROX (Invitrogen) on a ViiA7 qPCR detection system (Life Technologies). All reactions for RT-qPCR were performed using the following conditions: 50°C for 2 min, 95°C for 2 min, 40 cycles of a two-step reaction of denaturation at 95°C for 15 min and annealing at 60°C for 30s. The mean Ct values from triplicates were used in the comparative 2^-ΔΔCt^ method. To analyze the expression of Notch ligands on the adult muscle fiber, 2^-ΔCt^ method was used. The results were normalized to GAPDH mRNA controls. The primers used in this study are listed in Supplemental Table S2.

### Microarray

Total RNA was isolated from single muscle fibers of the EDL muscle from post-natal day 3 (p3), post-natal day 7 (p7) and adult mouse hindlimb using TRIzol reagent (Invitrogen). The processing of RNA, hybridization and analysis is previously described (Eliazer et al., 2019). The expression values for Notch ligands are log2 transformed and listed in Table S1.

## QUANTIFICATION AND STATISTICAL ANALYSIS

The density maps for Pax7 and Ddx6 intensity was drawn using R. The Variance of Pax7 and Ddx6 intensity levels across the SC pool are calculated as the square of standard deviation. The statistical details of experiments can be found in the figure legends. No statistical methods were used to predetermine sample size. The investigators were not blinded to allocation during experiments and outcome assessment. No animal was excluded from analysis. All data are represented as mean ± standard error of the mean (s.e.m). Significance was calculated using the two-tailed unpaired Student’s t-tests (Graphpad Prism 7). The number of replicates (n) for each experiment is indicated in the figure legends. Differences were considered statistically different at P<0.05.

## DATA AND CODE AVAILABILITY

The microarray data generated during this study is available at NCBI GEO: GSE135163

